# Development of an MR-guided focused ultrasound (MRgFUS) lesioning approach for small and deep structures in the rat brain

**DOI:** 10.1101/2023.10.11.561930

**Authors:** Carena Cornelssen, Allison Payne, Dennis Parker, Matthew Alexander, Robb Merrill, Sharayu Senthilkumar, Jacob Christensen, Karen S. Wilcox, Henrik Odéen, John D. Rolston

## Abstract

**Objective:** High-intensity magnetic resonance-guided focused ultrasound (MRgFUS) is a noninvasive therapy to lesion brain tissue, used clinically in patients and preclinically in several animal models. Challenges with focused ablation in rodent brains can include skull and near-field heating and accurately targeting small and deep brain structures. We overcame these challenges by creating a novel method consisting of a craniectomy skull preparation, a high-frequency transducer (3 MHz) with a small ultrasound focal spot, a transducer positioning system with an added manual adjustment of ∼0.1 mm targeting accuracy, and MR acoustic radiation force imaging for confirmation of focal spot placement.

**Methods:** The study consisted of two main parts. First, two skull preparation approaches were compared. A skull thinning approach (n=7 lesions) was compared to a craniectomy approach (n=22 lesions), which confirmed a craniectomy was necessary to decrease skull and near-field heating. Second, the two transducer positioning systems were compared with the fornix chosen as a subcortical ablation target. We evaluated the accuracy of targeting using a high-frequency transducer with a small ultrasound focal spot and MR acoustic radiation force imaging.

**Results:** Comparing a motorized adjustment system (∼1 mm precision, n=17 lesions) to the motorized system with an added micromanipulator (∼0.1 mm precision, n=14 lesions), we saw an increase in the accuracy of targeting the fornix by 133%. The described work allows for repeatable and accurate targeting of small and deep structures in the rodent brain, such as the fornix, enabling the investigation of neurological disorders in chronic disease models.

## Introduction

Transcranial high-intensity focused ultrasound (FUS) is a modern noninvasive therapy that uses thermal energy to lesion brain tissue (1). Additionally, magnetic resonance-guided FUS (MRgFUS) combines the superior soft tissue contrast of magnetic resonance imaging (MRI) with magnetic resonance thermometry imaging (MRTI) for real-time temperature monitoring (2,3). MRgFUS is approved by the United States Food and Drug Administration (FDA) to treat neurological disorders, such as essential tremor and Parkinson’s disease (2,3). FUS is advantageous over other modern noninvasive surgical techniques, such as stereotactic radiosurgery, as it does not use ionizing radiation and produces immediate results, enabling real-time feedback from awake patients (1,4). It also improves upon minimally invasive techniques like radiofrequency thermocoagulation and laser interstitial thermotherapy that involve passing a probe through normal brain parenchyma via a burr hole drilled in the skull (4,5). MRgFUS requires no incision and ideally only affects the targeted tissue (5).

MRgFUS is an accepted therapy for movement disorders but is also being investigated for other neurological disorders. Recently, MRgFUS has been used in a Phase 1, open-label clinical trial for drug-resistant epilepsy targeting the anterior nucleus of the thalamus and a reduction in seizures was observed in the two patients in the study (6). While these results are promising, other targets in the circuit of Papez, such as the fornix, may offer alternative approaches with less risk of ablating adjacent tissue (2). Before attempting MRgFUS for alternative targets in human epilepsy, however, the investigation of MRgFUS for use in a preclinical animal model is needed. Preclinical research is vital to studying epilepsy as it enables studies of therapeutic mechanisms and efficacies within different epilepsy phenotypes and syndromes, some of which are impossible to study in humans (7,8). There are numerous well-validated animal models in rodents of seizure-induced and chronic disease states of epilepsy, making rodents the primary choice of study (9).

A challenge with transcranial FUS in rodents is their small and flat skulls, necessitating using limited-size aperture transducers (10). This results in high acoustic power density as the FUS traverses the skull and can result in substantial skull and brain surface heating (1,10). The study consisted of two parts. In the first part, we compared two skull preparation methods, skull thinning and craniectomy, to limit skull heating and achieve heating at the desired deep brain target. The craniectomy approach avoided issues with skull heating and was used in the second part of the study, where we compared two different transducer positioning systems. We compared using only a piezoelectric motor positioning system alone to one with an additional manual positioning system. The motorized adjustment alone had an accuracy of ∼1 mm, while the additional manual adjustment resulted in an accuracy of ∼0.1 mm. Acoustic radiation force imaging (ARFI) was also used to minimize heat deposition during the targeting step. The work described in this paper allows for repeatable and accurate targeting of small targets deep in the rodent brain, such as the fornix. It is the groundwork for future studies, e.g., MRgFUS efficacy, for treating various forms of epilepsy.

## Materials and Methods

### Animal surgery

The University of Utah’s Institutional Animal Care and Use Committee granted approval for all animal procedures. During a one-day survival surgery under isoflurane anesthesia (1-3%, Figure 1a), all animals (N=40, male Sprague-Dawley rats, 230-500 g, Charles River Laboratories) underwent the same first step outlining a rectangle of ∼9 mm x 14 mm, centered on bregma, to shave down the skull to ∼200-micron thickness using a hand drill (Dremel 2050-15 Stylo+) and a spherical-shaped bur (Carbide bur, length of cut 1/16-inch, length 1.5 inches, Widia Metal Removal). Throughout drilling, chilled sterile saline (Medline Sterile Saline, 0.9%) was continuously washed over the skull to prevent drying of, and to cool, the skull bone.

**Figure 1:**
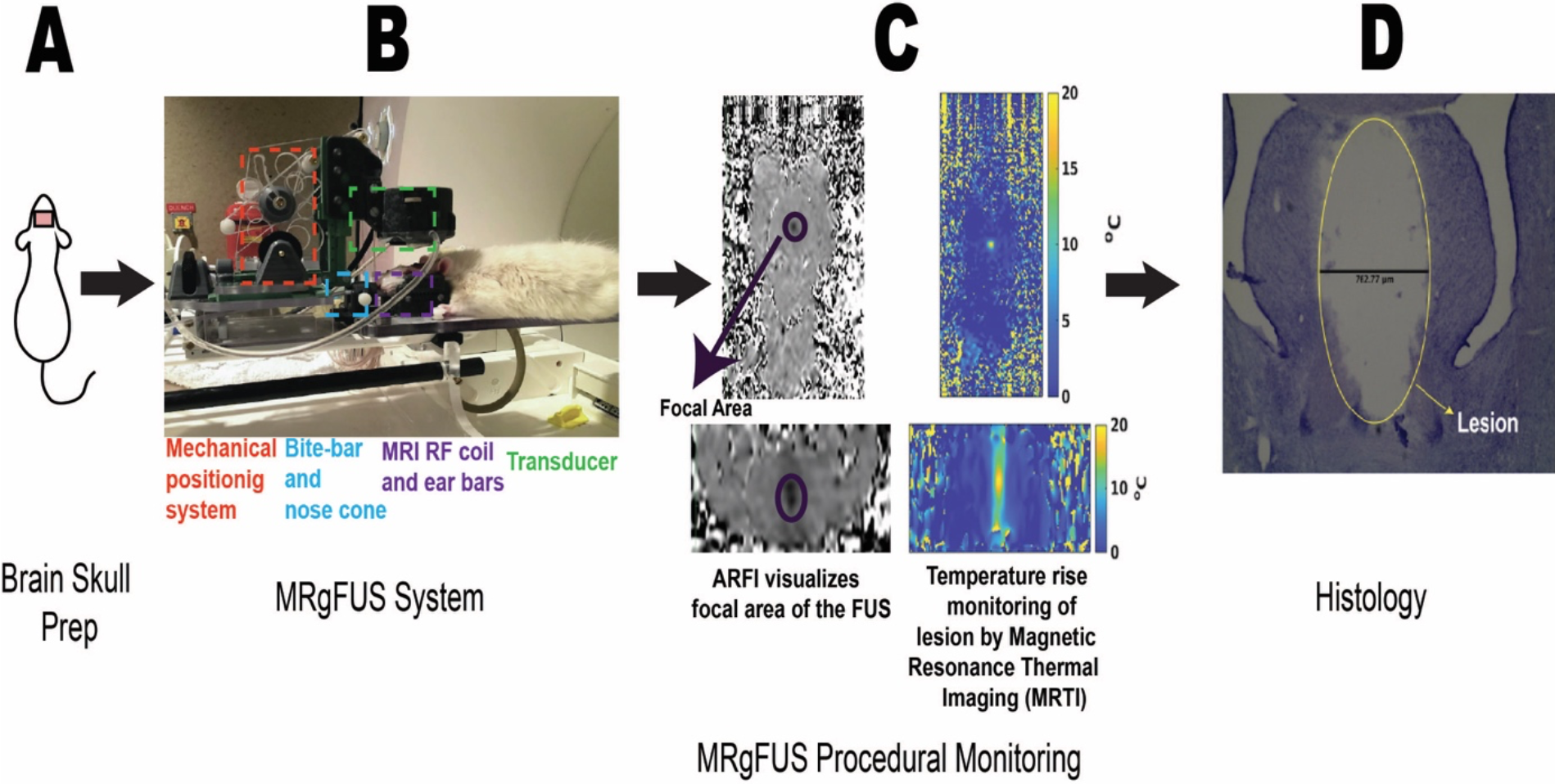
Experimental overview of the MRgFUS experiment. 1A shows the rat underwent a skull preparation method, which involved either skull thinning or a craniectomy. 1B, immediately following the skull preparation, the rat was positioned in a custom animal setup, including a bite bar, nose cone, and ear bars. A 3 MHz annular-array transducer was positioned above the skull opening. The in-house built mechanical adjustment, outlined in red, moves the transducer with sub-mm accuracy. 1C, once the rat was inside the MRI scanner, MRTI or ARFI was used to locate the focal area. MRTI was used to monitor the temperature rise as the thermal lesion was created. 1D, 24 hours or 5 days following the procedure, the rats were perfused, brains harvested and sliced, and slices underwent Cresyl Violet staining for verification of the accuracy of the lesion and tissue lesioned around the intended target, the fornix.

### Skull thinning preparation

In a subset of the rats (n=7, 230-400 g) shown in Table 1, the skull was shaved down to ∼50-microns thickness with a diamond cylinder bur (Dedeco® Goldies® HP Diamond Point 78/4060, titanium nitrate coated bur). The resulting thinning skull prep was characterized by being flexible to the touch and translucent.

**Table 1:**
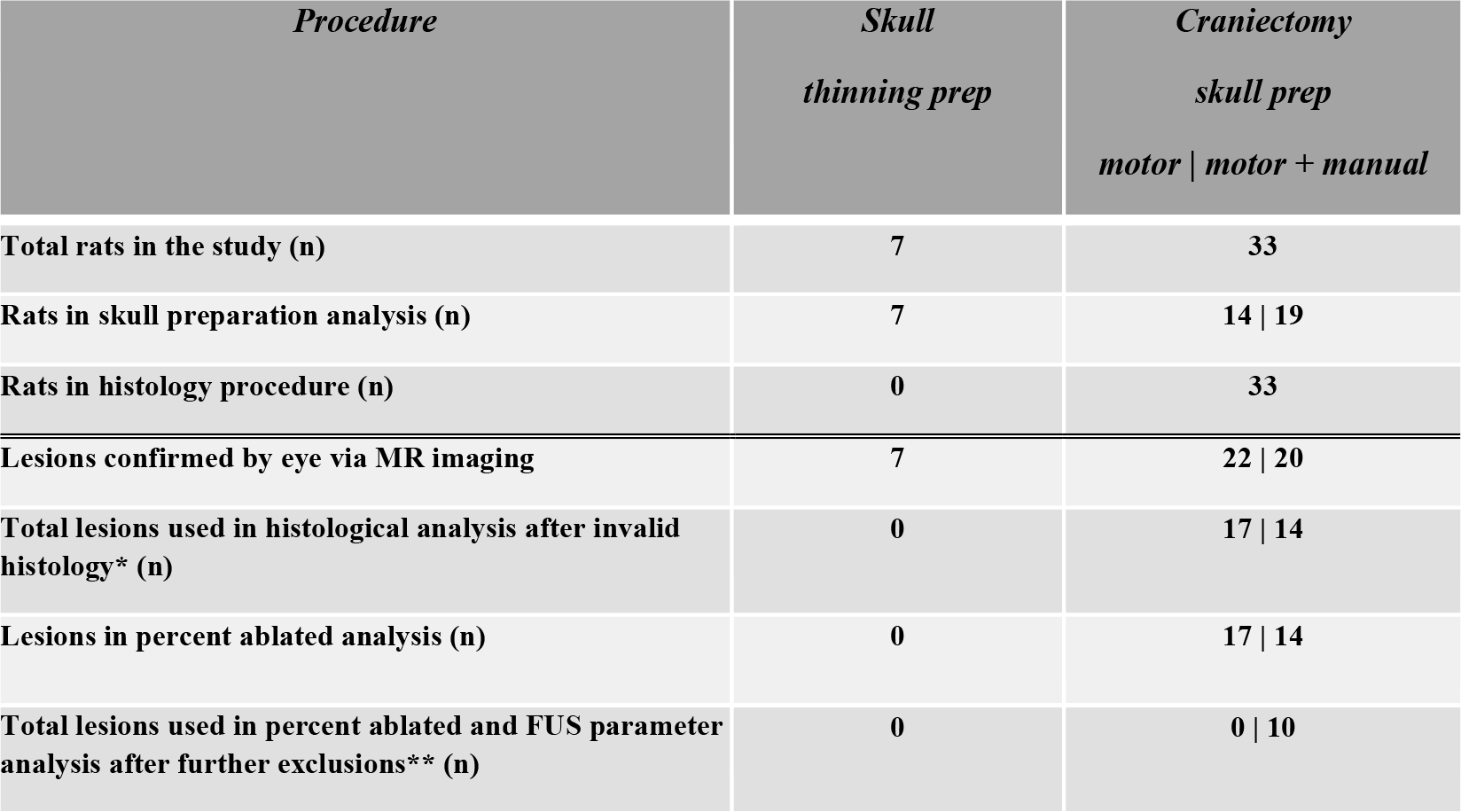
Number of rats and lesions across study procedures and analyses. Displays the number of rats and lesions (below the blue line) in various procedures and analyses in the study for the skull thinning and craniectomy skull preparation methods. Where applicable, numbers are broken down into the study’s two parts (motor and motor+manual). *Invalid histology reasons included target not seen in histology, or the tissue was unusable. **Further exclusions were due to multiple sonications and/or lesions on the same brain, which would exclude that lesion from being comparable to lesions with a single sonication for analysis.

| *Procedure* | *Skull*<br>*thinning prep* | *Craniectomy*<br>*skull prep*<br>*motor \| motor + manual* |
| --- | --- | --- |
| Total rats in the study (n) | 7 | 33 |
| Rats in skull preparation analysis (n) | 7 | 14 \| 19 |
| Rats in histology procedure (n) | 0 | 33 |
| Lesions confirmed by eye via MR imaging | 7 | 22 \| 20 |
| Total lesions used in histological analysis after invalid histology* (n) | 0 | 17 \| 14 |
| Lesions in percent ablated analysis (n) | 0 | 17 \| 14 |
| Total lesions used in percent ablated and FUS parameter analysis after further exclusions** (n) | 0 | 0 \| 10 |

### Craniectomy

In another subset of the rats (n=33, 230-500 g), the skulls were shaved along the outline of the rectangle until the dura mater was reached, and fine forceps were used to remove the rectangular piece of the skull. Prior to the surgery and use, hemosponges (Goodwill Hemosponge Premium Quality, Dental World Official) were cut into even pieces and degassed in sterile saline (<2 ppm dissolved oxygen) for one hour. The hemosponges were placed directly on the craniectomy site on the rat’s head to achieve hemostasis and form a barrier between the brain and the external environment.

### MRgFUS procedure

Directly after the craniectomy (Figure 1A), the rats were placed in our MRgFUS apparatus (Image Guided Therapy, IGT, and Imasonic), which was specifically designed for small animals (Figure 1B). Our setup consists of a 16-channel annular array 3 MHz transducer (Imasonic) with a focal spot size of 1x1x3 mm (full width at half maximum, free-field measured by a hydrophone in a water bath) capable of electronically steering in the ultrasound propagation direction, a custom receive-only MRI radiofrequency (RF) coil built in-house, which allows for positioning close to the animal for increased image SNR, ear- and bite-bars which are used to position and secure the animal’s head, and a nose cone to deliver inhalational anesthesia during the procedure. An enlarged figure detailing the design and characteristics of the MRgFUS positioning setup from Figure 1B is shown in Figure 2.

**Figure 2:**
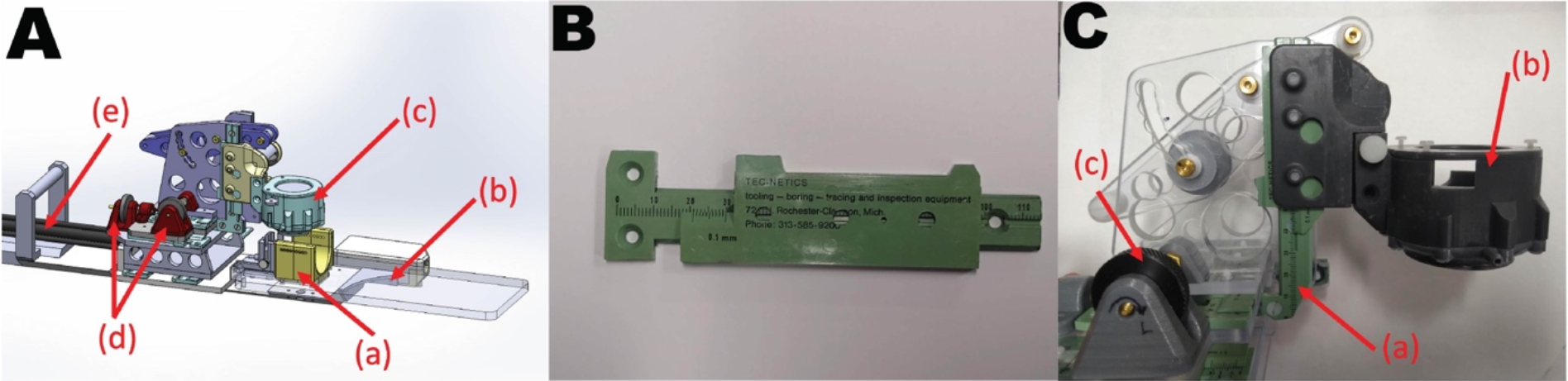
MRgFUS positioning setup. A, CAD drawing of the design showing (a) head-holder with built in RF-receive coil and holes for ear bars, (b) animal support table, (c) housing for 3-MHz transducer, (d) knobs for manual position adjustment, (e) shafts going to piezoelectric MR compatible motors. B, the manual adjustments in x, y, and z were designed and built using modified MR-compatible plastic calipers (Fowler, Mausner Equipment Company). The precision of the adjustments was 0.1 mm, with x-y (head-foot and right-left, respectively) ranges of 7 mm and z (anterior-posterior, i.e., up-down) range of 40 mm. C, close-up photo of part of the finished design showing (a) calipers, (b) transducer housing, and (c) one of the knobs for manual adjustments (x, for head-foot adjustments).

Rats were placed prone, and their heads secured with a combination of the nose cone and ear- and bite-bars in the MRI (3T, PrismaFIT, Siemens Healthineers) under continued isoflurane anesthesia (1-3%, Figure 1B). The FUS transducer was coupled to the animal’s head by a deionized and a degassed water bath contained by a flexible membrane and water-based sterile acoustic coupling gel (Aquasonic 100). The skin on the head was held back from the FUS coupling cone with small, loose, removable stitches (Nylon, 5-0, Non-Absorbable, FS-2, 18”, Ethicon) on both sides of the head.

High-resolution MR imaging was performed to identify the fornix and ablation regions. T2-weighted 3D Sampling Perfection with Application optimized Contrast using different flip angle Evolution (3D SPACE, field of view = 134 x 50 x 28 mm, 0.35 x 0.35 x 0.5 mm voxel size, reconstructed at 0.17 x 0.17 x 0.25 mm voxel spacing, TR = 1450 ms, TE = 82 ms, 2 averages, BW = 435 Hz/px, acquisition time = 6 min 41 s) was used. The fornix was identified adjacent to the anterior commissure (AC)-posterior commissure (PC) line in the sagittal plane of the anterior commissure. The target was assigned at or just posterior to the AC–PC line. Moving along the midline in the scan sequence when the AC is in view, but the cingulate gyrus is not, we assigned the fornix target just posterior to the AC and bordering the third ventricle. Here, the ablation occurred, shown by the target in the red arrow in Figure 3.

**Figure 3:**
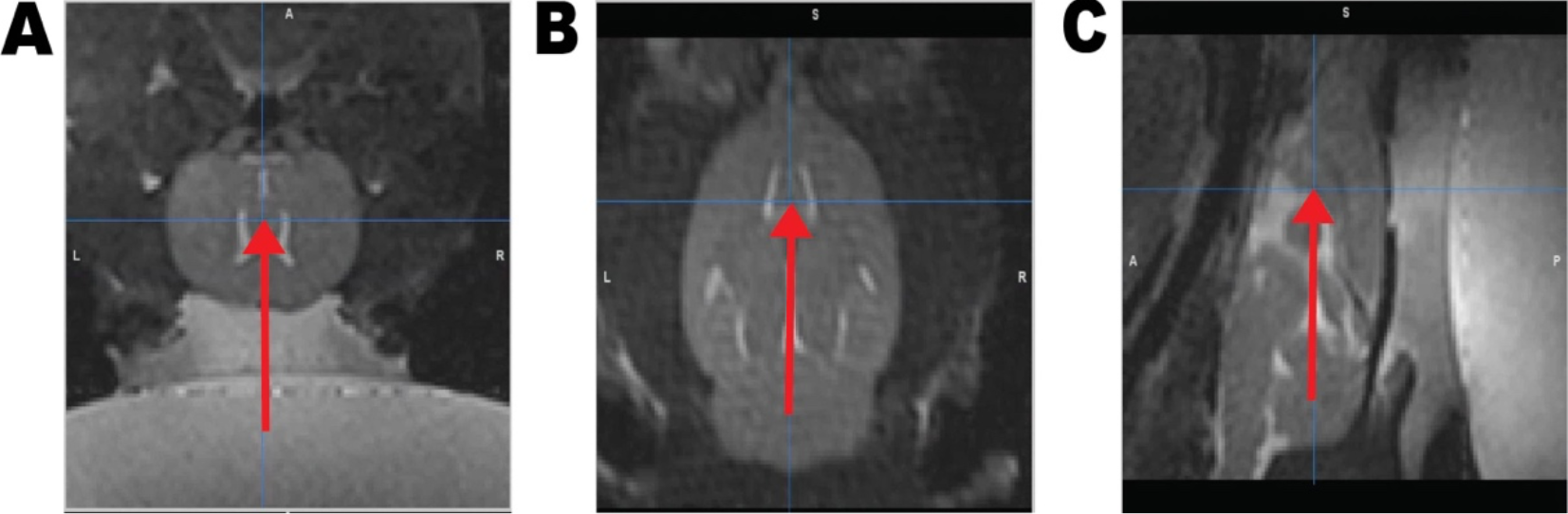
Target for ablation. The fornix target for ablation is located at the end of the red arrow in a coronal view in 3A, axial/transverse view in 3B, and sagittal view in 3C using a T2w SPACE sequence at 0.35 x 0.35 x 0.5 mm resolution.

In the first part of the study, target verification was performed using low-power sonications monitored by MRTI (3D segmented echo planar imaging (EPI), field of view=96 x 48 x 18 mm, 1.5 mm isotropic voxel size, reconstructed at 0.75 mm isotropic voxel spacing, TR = 25 ms, TE = 15 ms, BW = 744 Hz/px, echo train length=3, 2.5 s/3D volume), and in the second part of the study, MR-ARFI (3D segmented EPI, field of view=96 x 48 x 18 mm, 0.6 x 0.6 x 1.5 mm voxel size, reconstructed at 0.3 x 0.3 x 0.75 mm voxel spacing, TR = 50 ms, TE = 27 ms, BW = 744 Hz/px, echo train length=3, motion encoding gradients 7.5 ms, 50 mT/m, 24 s/ARFI displacement image) was used to visualize the focal spot before creating a permanent lesion. Our custom MRgFUS system was used to create lesions of the fornix (-0.60 mm A/P, 0 mm M/L, 5.0 mm D/V). Pre- and post-ablation MRI (T2-weighted SPACE, Figure 3) was performed to identify and evaluate acute lesion size and location. MRTI was performed and visualized in real-time (Thermoguide™, IGT) for monitoring the treatment (Figure 1C). Sonication duration varied between 4.95 – 10.00 seconds, depositing between 3.67 – 64.22 J resulting in a maximum temperature of 40 – 67 °C at the target, measured with MRTI. Both single and multiple sonications in the same spot were used to induce lesions.

### Skull preparation analysis

The first part of the study compared lesions via MR imaging for the skull thinning preparation (n=7) to the craniectomy with the first transducer system (n=22), shown in Table 1. The overlap of the intended target and the heating location was compared through visual inspection of MRTI overlaid on the post-treatment T2w MRI (Figure 4). This part of the study used only the MR-compatible motors (i.e., not the manual adjustment system) for transducer positioning. The temperature scale indicates the absolute temperature (°C), with red areas having a higher temperature, indicating potentially ablative temperatures of > 55°C (1,10).

**Figure 4:**
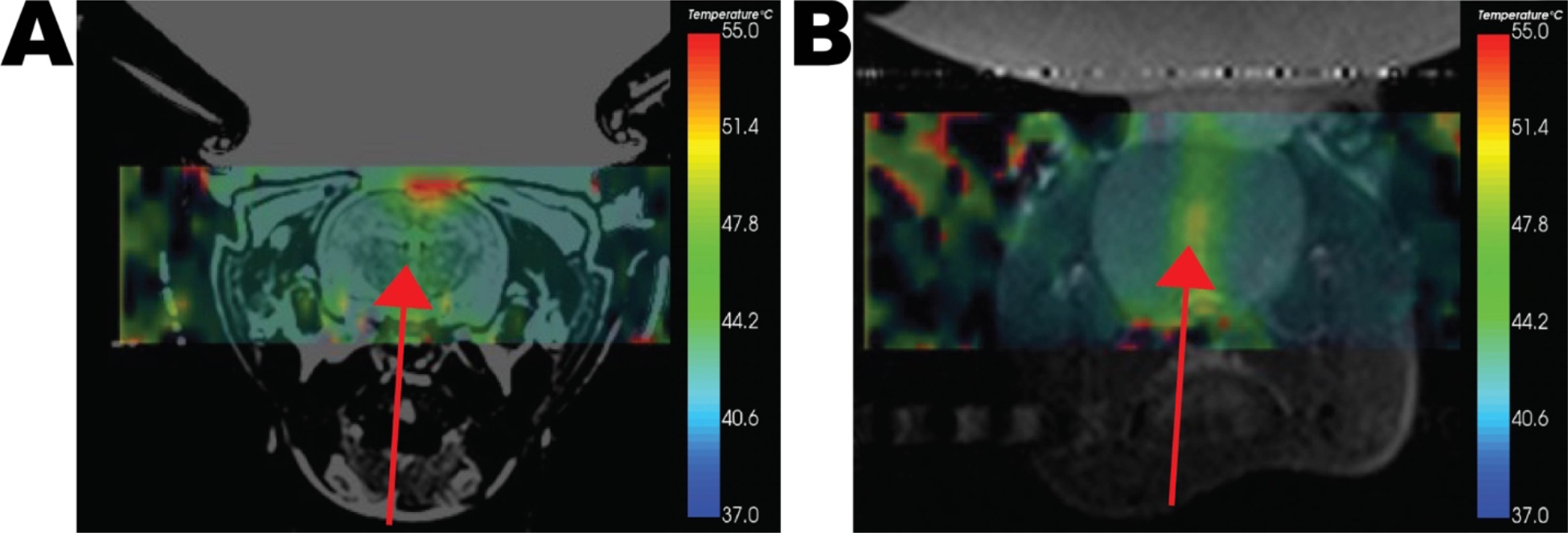
Comparison of skull preparation methods. Representative figures comparing the skull thinning prep (A) and craniectomy (B) methods visually through MRTI overlaid on the post-treatment T2w. The red arrow displays the intended target. 4A, the rat underwent a skull thinning prep, resulting in substantial near-field heating reaching around 55°C. 4B, the rat underwent a craniectomy. At the end of the red arrow is a red heat area representing the ablated area, resulting in no heating in the ultrasound focal near-field, but heating around 55°C at the target. Thus, the craniectomy preparation method was superior to the skull thinning prep method.

### Histology

All rats that underwent the craniectomy (n=33) were used for the histological analysis, shown in Table 1. Twenty-six animals underwent transcardiac perfusion within 24 hours of the MRgFUS, and the remaining seven underwent perfusion by day five, using 4% paraformaldehyde (Thomas Scientific) in 1x phosphate buffer saline (Dulbecco’s). Following removal, the brains were kept in 4% paraformaldehyde for 48 hours, transferred to 30% sucrose for 96 hours, and then sectioned on a microtome (SM 2010 R, Leica) at a 90-micron thickness. The lesion size and location created by the MRgFUS fornicotomies were verified through histology using a Cresyl Violet stain to evaluate structural changes to the tissue (Figure 1c) (11).

### Histological analyses

After excluding 11 lesions due to invalid histology (e.g., lesion was not observed in tissue, or the tissue was unusable), 31 lesions were analyzed. We compared the motorized adjustment lesions (n=17) to the added manual adjustment lesions (n=14) for the initial comparison between the transducer positioning systems, shown in Table 1. The lesions were viewed under brightfield microscopy (XL Core, EVOS) at 40x to determine the accuracy of the lesion to the target, the fornix (-0.60 mm A/P, 0 mm M/L, 5.0 mm D/V), as well as the neighboring structures lesioned, using a standard rat atlas during visual inspection of lesions (12).

After the determination that the motorized+manual positioning system provided superior accuracy, a sub-analysis was performed with these lesions (n=14) investigating the percent volume of the fornix and surrounding structures that were ablated and what effect FUS parameters (temperature rise (°C) and total energy deposited (J)) had on the ablation volume. We controlled for the number of sonications that made a lesion, which could affect the lesion size and structures ablated. Thus, an additional N=4 lesions from the N=14 sub-analysis group were excluded for a total of 10 lesions to determine which FUS parameters optimized lesion size shown in Table 1. MATLAB (2022b) was used to plot and display the results in all analyses.

### Accuracy of targeting the fornix analysis

Accuracy was evaluated by comparing where the lesion occurred with respect to the intended target. All lesions (n =31), whether a full ablation of the target or not, were included in this analysis to study the improvements made to the system setup. Lesions were grouped for accuracy as either on target, anterior to the target, posterior to the target, or missing the target. ‘On target’ means the fornix was transected at the level targeted. ‘Anterior’ and ‘posterior’ to the target means a lesion that partially or fully transected the fornix, however not at the coordinate intended. ‘Miss’ means a lesion that did not hit any component of the fornix. Accuracy of being on target was considered transecting the fornix at the coordinate intended (-0.6 mm A/P, 0 mm M/L, 5.0 mm D/V) (12).

### Percent ablated analysis

After visual inspection of all cross-sections of the lesion under the microscope, any structure that was at least partially ablated was included in the analysis. The percent ablated was calculated using the volume of the anatomical structure.

The height and width of the ablated volume of a structure were determined by measuring lesions in the cross-sectional histology slices using ImageJ (version 1.53a for Mac). To avoid bias, as the structure may not always occur in the largest cross-section of the lesion, we took measurements from the cross-section that had the largest area ablated of the structure to use when calculating volume ablated for that structure. The first and last cross-sections of the anatomical structure ablated were matched to the standard rat atlas to determine the respective bregma coordinates (12). The distance from the first and last cross-sections was measured to determine the length of the ablated volume by subtracting the coordinates. The lesion volume in each structure was estimated using the formula for an ellipse, as the lesion was assumed to be the shape of an ellipse based on the cigar-shape of the ultrasound focus. While most structures require ablating the full anatomical structure to cease their functions, white matter tracts, such as the fornix, require ablating only a cross-section (transection). Therefore, any white matter tract structure was considered to have an ablated structure volume of 100% (fully ablated) when an entire cross-section was ablated.

To compute the average volume of a structure, we used already validated mean brain volumes for the following structures: hippocampus, corpus callosum, cingulate cortex, sensorimotor cortex, thalamus, internal capsule, and caudate putamen from Welniak-Kaminska et al. (13). When structures were unavailable in Welniak-Kaminska et al., we calculated the standard structure volume from average measurements from the standard rat atlas (12). Percent ablated was then calculated by dividing the measured ablation volume by the average structure volume for the same structure.

### FUS parameter analysis

The same lesions used for the percent ablated analysis were used to investigate which FUS parameters (power and duration) could repeatedly produce a lesion size similar to the fornix of 0.68 mm^3^, 2 x 1.3 x 0.54 mm in volume and height x width x length, respectively (12). All lesions were used to determine if a correlation existed between lesion volume and temperature rise, and energy deposited. A linear regression was completed for the FUS with lesion volume (mm^3^) to determine the correlation of each parameter to lesion size. All data analysis was performed in MATLAB (2022b).

## Results

We compared two skull preparation methods (a thinning skull preparation and a craniectomy) and two transducer positioning systems and determined the accuracy of ablating the fornix and lesion size. Additionally, in a sub-analysis using added manual adjustment lesions, we investigated the percent of volume of both the target that was ablated and any off-target structures, as well as determine which FUS parameters affect lesion volume.

### Compared to the skull thinning preparation method, the craniectomy results in less near-field heating

Lesion locations were visually compared by looking at the MRTI overlaid on post-treatment T2w to compare temperatures of 55°C known to cause ablation in rats of the skull thinning method to the craniectomy method (Figure 4) (10). Through visual comparison of lesion areas compared to the intended target at the outlined red arrow (Figure 4), we confirmed that the craniectomy approach (Figure 4B, n=17) yielded heating at the target shown in red to be 55°C through the temperature scale bars. The craniectomy approach also yielded less skull and near-field heating shown by lack of 55°C heating in those areas when compared to the skull thinning prep method (n=7; Figure 4A).

### The addition of a manual adjustment increased the accuracy of targeting

Figure 5 shows that with the added micromanipulator, the accuracy of targeting the fornix increased by 133% and decreased misses by 100%, as compared to only using a motorized adjustment.

**Figure 5:**
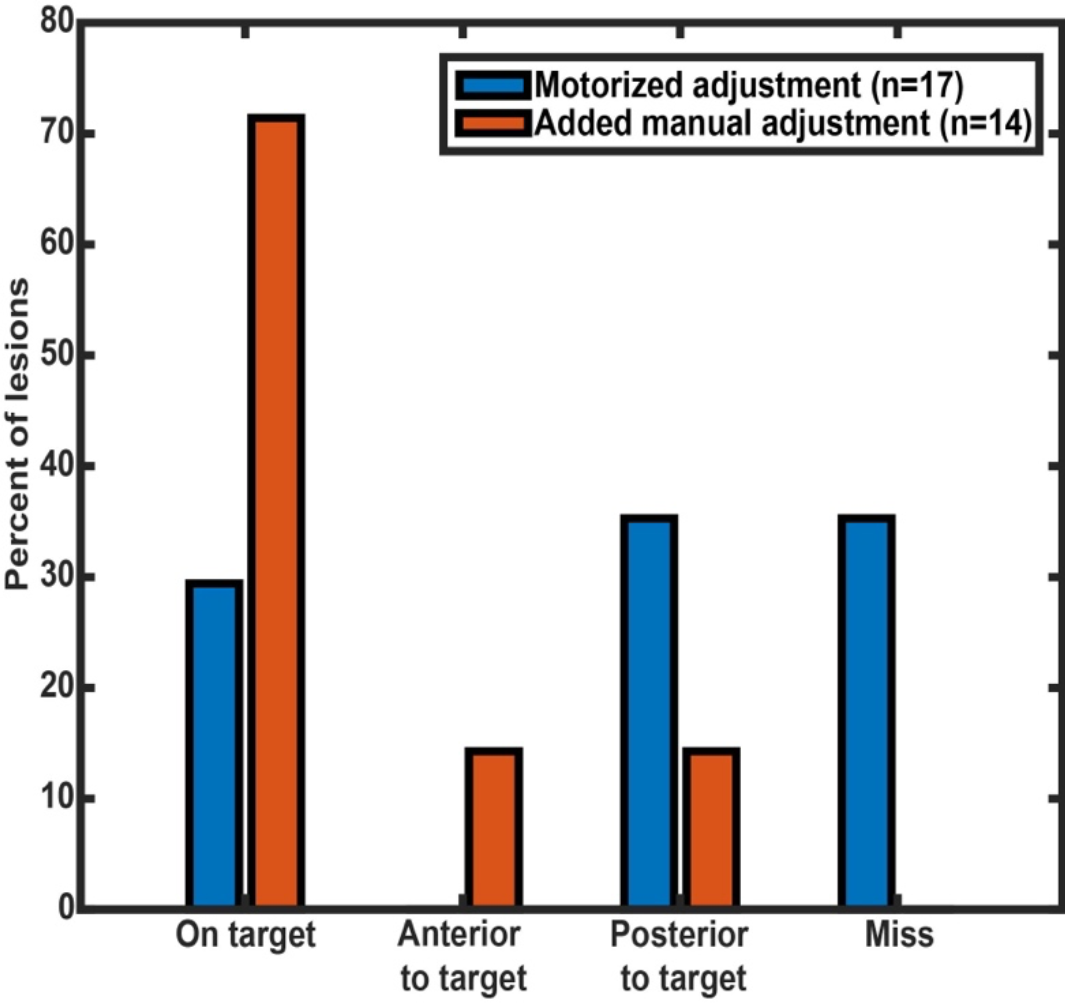
Accuracy of lesioning the fornix comparing two transducer positioning systems (n =31). Ablating the fornix at the coordinates (-0.60 mm A/P, 0 mm M/L, 5.0 mm D/V) was ablating ‘on target,’ ablating the fornix anterior to the coordinates was considered ‘anterior to the target’ and ablating posterior to the coordinates was considered ‘posterior to the target.’ Whereas ‘miss’ completely missed the target.

### Greater ablation was observed in the fornix when the manual adjustment was added

Following the addition of the manual manipulator, the majority of the ablation occurred within the fornix (Table 2), and less than 10% of the neighboring tissues were ablated (Table 3). Investigating the added manual adjustment lesions, we achieved complete ablation in 40% of our lesions. In 10% of our lesions, we ablated 85% of our target. In 50% of the lesions, we ablated ∼20% of the target.

**Table 2:** Percent of the target (fornix) ablated. The percent of volume ablated of the target structure (fornix) when using the manual lesion adjustment for each lesion (n=10).

| Lesion ID | Percent ablated of the fornix |
| --- | --- |
| 1 | 100.0 |
| 2 | 100.0 |
| 3 | 85.1 |
| 4 | 99.6 |
| 5 | 100.0 |
| 6 | 23.7 |
| 7 | 14.6 |
| 8 | 16.2 |
| 9 | 31.3 |
| 10 | 8.9 |

**Table 3:** Average percent ablated of the fornix and off-target structures. The percent of volume ablated at the on- and off-target structures (neighboring structures to the fornix) when using the manual lesion adjustment for each lesion using sub-analysis lesions (n=10).

| Structure | Number of lesions with the structure ablated | Average percent ablated |
| --- | --- | --- |
| fornix | 10 | 58 |
| hippocampus | 1 | 0.0 |
| corpus callosum | 6 | 0.1 |
| sensorimotor cortex | 1 | 0.0 |
| ventral hippocampal commissure | 1 | 3.5 |
| thalamus | 1 | 0.0 |
| triangular septal nucleus | 1 | 2.6 |
| stria medullaris | 0 | 0.0 |
| subfornical organ | 2 | 9.5 |
| retrosplenial agranular cortex | 0 | 0.0 |
| medial habenular nucleus | 0 | 0.0 |
| internal capsule | 0 | 0.0 |
| stria terminalis | 0 | 0.0 |
| caudate putamen | 0 | 0.0 |
| lateral septal nuclei | 10 | 0.2 |
| medial lemniscus | 0 | 0.0 |

### FUS parameters showed a significant correlation to lesion size

We compared the FUS parameters in the added manual adjustment (n=10) to understand how to optimize lesion size while increasing ablation at the fornix and decreasing ablation at the neighboring structures. When performing a linear regression analysis, both the temperature rise (°C) and energy deposited (J) FUS parameters showed a positive correlation with the volume of the lesion (Figure 6).

**Figure 6:**
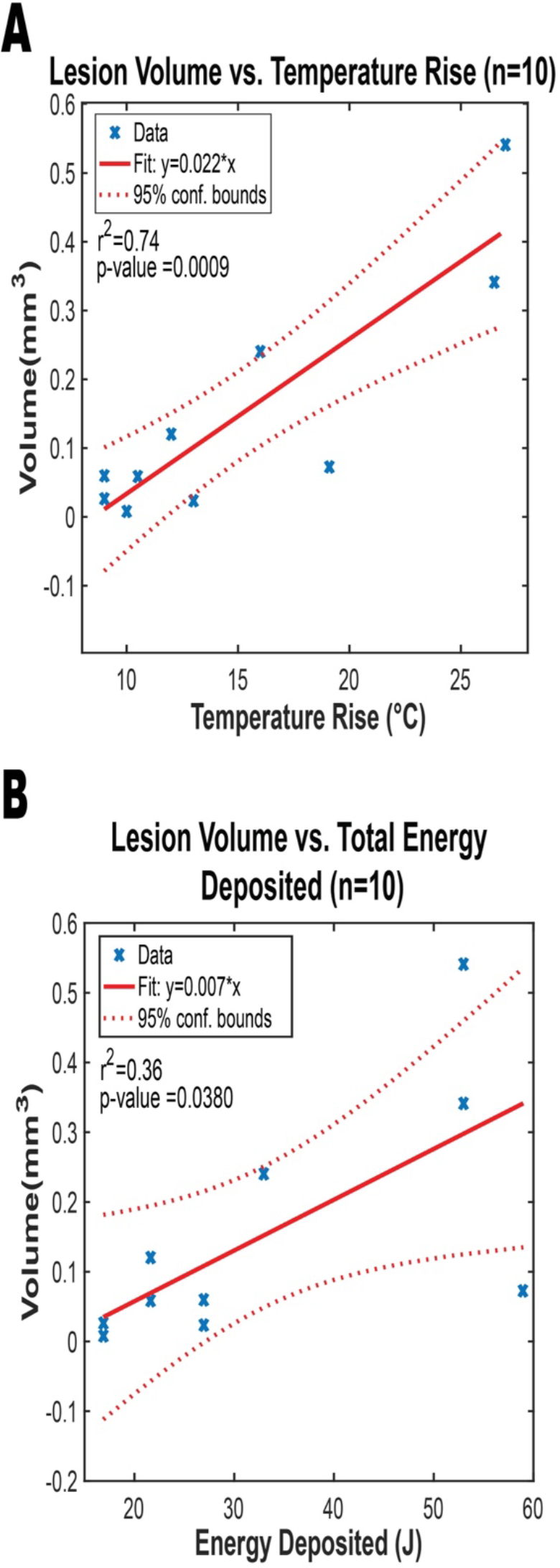
Comparison of the FUS parameters to the lesion size. An increase in the change in temperature (A) and an increase in energy deposited (J) (B) indicated an increase in lesion volume.

## Discussion and Conclusion

Prior to human application, small animals, especially chronic disease models, are used to investigate the effects of therapies. However, in the field of FUS, there has been an obstacle when using small animals: the geometry of the animal’s skull has a different shape than the human head, which necessitates using a different aperture-size transducer. A common problem when small aperture-size transducers are used is that the incident acoustic energy is spread over a smaller area, resulting in increased skull heating. In the first part of the study, we documented and overcame this obstacle by showing the superiority of a limited craniectomy over skull thinning in adult rodents. To further refine targeting capabilities of small anatomical targets in the rodent brain, we developed a micropositioning system that could be coupled to our piezoelectric positioning system. The added manual adjustment increased precision by 133%, which is beneficial when targeting structures smaller than 1 mm in the rodent brain. When trying to achieve a specific lesion size, especially for small brain structures, we showed that both temperature rise and total energy deposited were important FUS parameters that contribute to lesion size. This can be useful when planning future studies or treatments.

The limitations of the study included, foremost, the necessity of a craniectomy. A craniectomy was needed to prevent skull burns and near-field heating. Craniectomy is unacceptable in humans undergoing MRgFUS, though. However, in humans, the target size is larger, and the skull is more favorable to heating central structures like the fornix, meaning that preclinical findings investigating certain targets in rodents (like the fornix) could be translated to humans using standard FUS approaches. This will still allow a comparison of targeted subcortical ablations without transgressing brain en route to these targets, as required with radiofrequency ablation, for example. An alternative to the craniectomy approach would be to investigate MRgFUS ablations in larger animals. Though the tradeoff there is the paucity of chronic disease models and costs of larger animals. Another approach would be the use of substantially younger rodents, where thinner skulls may allow ablation. However, many disease models take longer to develop, limiting the utility of that approach.

In conclusion, we developed an improved methodology for targeting and ablating small and deep brain structures in rodents. This may allow us to study and identify novel targets for MRgFUS trials in humans.

## Acknowledgments

This work was supported by the following grants, NSF Graduate Research Fellowship 1747505 (C.C.), Richard L. Stimson Endowed Chair (K.S.W.), Research Incentive Seed Grant, University of Utah School of Medicine (J.D.R.), NIH R01EB028316 (DLP), R21EB033638 (H.O.), and S10OD018482 (D.L.P.).

## Data Availability

Raw data files are available upon request.

## Contributions

J.D.R. conceived the study. C.C. created the study design and managed the study. C.C., A.P., H.O., and J.C. performed the experiments. M.A. created a targeting technique and assisted with experiments. R.M. designed and developed the small animal holder and manual adjuster. C.C. and J.C. performed histological preparations. C.C. and S.S. analyzed the results. D.P. and K.S.W. commented on the results and provided expertise on the study. C.C. wrote the manuscript. C.C., A.P., D.P., KSW, H.O., and J.D.R. revised the manuscript. All authors approved the final manuscript.

